# Saturating variant analysis of TERC redefines the human telomerase reverse transcription mechanism

**DOI:** 10.64898/2026.06.01.728939

**Authors:** William Mannherz, Luke Homfeldt, Noah Lampl, Suneet Agarwal

## Abstract

Genetic variation at the *TERC* locus, encoding the telomerase RNA component, is associated with human diseases and lifespan, but a comprehensive functional annotation of this critical non-coding RNA is lacking. Here, we performed saturating variant analysis of TERC in cells and unexpectedly redefine the human telomerase catalytic mechanism. Telomerase is understood to precisely reverse transcribe six TERC templating residues to generate GGTTAG repeats, however our screen identified dominant negative effects expected of templating bases inconsistent with this annotation. In vitro and in cells, we found human telomerase uses not six, but eight TERC residues flexibly for reverse transcription, most commonly yielding GGGTTA, but variably up to AGGGTTAG. An evolutionary change in TERC adjacent to the template explains the long-standing misannotation, reversion of which shifts the main templating register in TERC back to the one assumed for decades. Remarkably, this template-adjacent change also yields hyperactive TERC variants that rapidly lengthen telomeres when introduced into cells, including those from patients with genetic telomere diseases. By functional analysis of *TERC* variation in human cells, our work revises core tenets of telomerase reverse transcription and provides a new mechanistic model to inform the development of telomere-directed therapeutics.

---

Telomere maintenance is critical in human health and disease. During genome replication, the terminal portion of telomeres is lost because DNA polymerases cannot fully replicate linear chromosomes^1–3^. Telomere length can be replenished by telomerase, an unusual reverse transcriptase holoenzyme complex that in humans includes the catalytic subunit TERT and an intrinsic non-coding RNA component called TERC^4–8^. TERC was cloned over three decades ago based on predictions of its template domain sequence^6^, followed by extensive functional, structural and phylogenetic studies revealing RNA domains and biogenesis features critical for telomerase function in vitro and in cells^9–12^. Recent cryo-EM studies validate and extend years of biochemical studies indicating that TERC scaffolds two functionally separable lobes of the telomerase holoenzyme: 1) the TERC template, pseudoknot and CR4/5 domains complexed with TERT, which is sufficient to reconstitute reverse transcriptase activity in vitro, and 2) the TERC hairpin-hinge-hairpin-tail structure complexed with the dyskerin H/ACA ribonucleoprotein and TCAB1, required for accumulation and trafficking of telomerase in human cells^13–15^. In the human population, genetic variation at the *TERC* locus shows the highest genome-wide association with telomere length, which in turn is associated with longevity and degenerative disorders^16,17^. In Mendelian telomere biology disorders (TBDs)^18,19^, mutations in *TERC* and associated factors cause disease by diverse molecular mechanisms, including defects in TERC biogenesis, stability, template function, assembly of the dyskerin H/ACA RNP, subcellular trafficking, and association with TERT^20^. Collectively these studies have provided a detailed mechanistic understanding of how telomerase maintains genome integrity and cellular self-renewal capacity throughout the lifespan and across generations, and how this process goes awry in degenerative diseases, aging, and cancer.

The canonical view of the telomerase catalytic cycle is that the telomere end hybridizes to complementary nucleotides at TERC positions n.56-52 and is extended by reverse transcription of the sequence 5’-GGTTAG-3’ precisely templated by TERC residues n.51-46^21,22^. The newly extended telomere end then dissociates and re-hybridizes to TERC alignment positions n.56-52, and the reverse transcription cycle is repeated to achieve processive synthesis of tandem 5’-GGTTAG-3’ DNA repeats. Both the telomere repeat and the templating residues of TERC are highly conserved – a common hexanucleotide telomere sequence 5’-TTAGGG-3’ is found across diverse eukaryotes from filamentous fungi to humans, suggesting it to be the common ancestor from which certain other telomere repeat sequences diverged^23,24^. Telomere repeats are bound with high sequence specificity by the shelterin complex of proteins^25,26^, and the single-stranded G-rich overhang of the telomere hybridizes internally to form the t-loop^27^. Together these structures shield the free chromosome terminus and promote genome integrity by suppressing DNA damage signaling and preventing end-to-end fusions^28^.

The alignment and template annotations of TERC have been undisputed for decades. The functional importance of other residues across the 451 nucleotide TERC RNA sequence has not been fully defined. It is clear that genetic variants spanning the TERC coding region impair telomerase function via various mechanisms to cause a spectrum of fatal genetic TBDs. In clinical practice, variants of uncertain significance in TERC are increasingly identified and pose diagnostic and therapeutic challenges, especially for TERC as a non-coding RNA where most *in silico* pathogenicity tools cannot be applied. We thus performed saturating variant analysis of the *TERC* gene, examining all possible single nucleotide changes in the primary sequence for their impacts on telomerase function in human cells, with the goal of cataloging and better understanding genetic mutations that contribute to TBD pathogenesis. Unexpectedly, our studies reveal novel mechanisms of human telomerase reverse transcription, including reassignment of core TERC templating residues designated 30 years ago.

## Saturating variant analysis of human TERC reveals genetic vulnerabilities

To enable high throughput functional screening of TERC variants (**Fig. 1a**), we generated a lentiviral vector which at low multiplicity of infection drove TERC expression ∼6x endogenous levels and yielded robust telomere elongation in TERC-null human embryonic kidney 293T cells (Extended Data Fig. 1a-c)^29^. We generated all possible 1353 single nt variants of the 451 nt TERC sequence in this vector, split across five sub-libraries with representation confirmed by next generation sequencing (NGS) (Extended Data Fig. 1d-e). TERC-null 293T cells divide normally until telomeres undergo replication-associated attrition to a critically short length, at which point cells senesce and stop growing (**Fig. 1a**)^29,30^. We introduced the saturating variant library into TERC-null 293T cells with the hypothesis that TERC variants that significantly impair or alter its function are unable to rescue these cells from senescence and would be depleted from culture. Following lentiviral infection, we cultured cells infected with the TERC variant libraries for 40 days alongside uninfected and WT TERC vector-infected controls, at which point the uninfected cells had stopped dividing (**Fig. 1b**). We isolated genomic DNA from cells infected with the TERC libraries, PCR amplified TERC transgenes from integrated lentiviruses, performed NGS, and quantified relative variant abundance compared to the input plasmid libraries^31^. We found depletion of variants across regions known to be essential for TERC function and accumulation, including the template domain, pseudoknot, CR4/5 domain, and the box H/ACA domain (**Fig. 1c,d**, Extended Data Fig. 2a-e)^10,32^. At the single nucleotide level, we found a strong correlation between evolutionary conservation of residues and their depletion in our screen (Extended Data Fig. 2f)^33^. Together, these data validate our screening approach to identify sequences critical for TERC function in human cells.

**Fig. 1.**
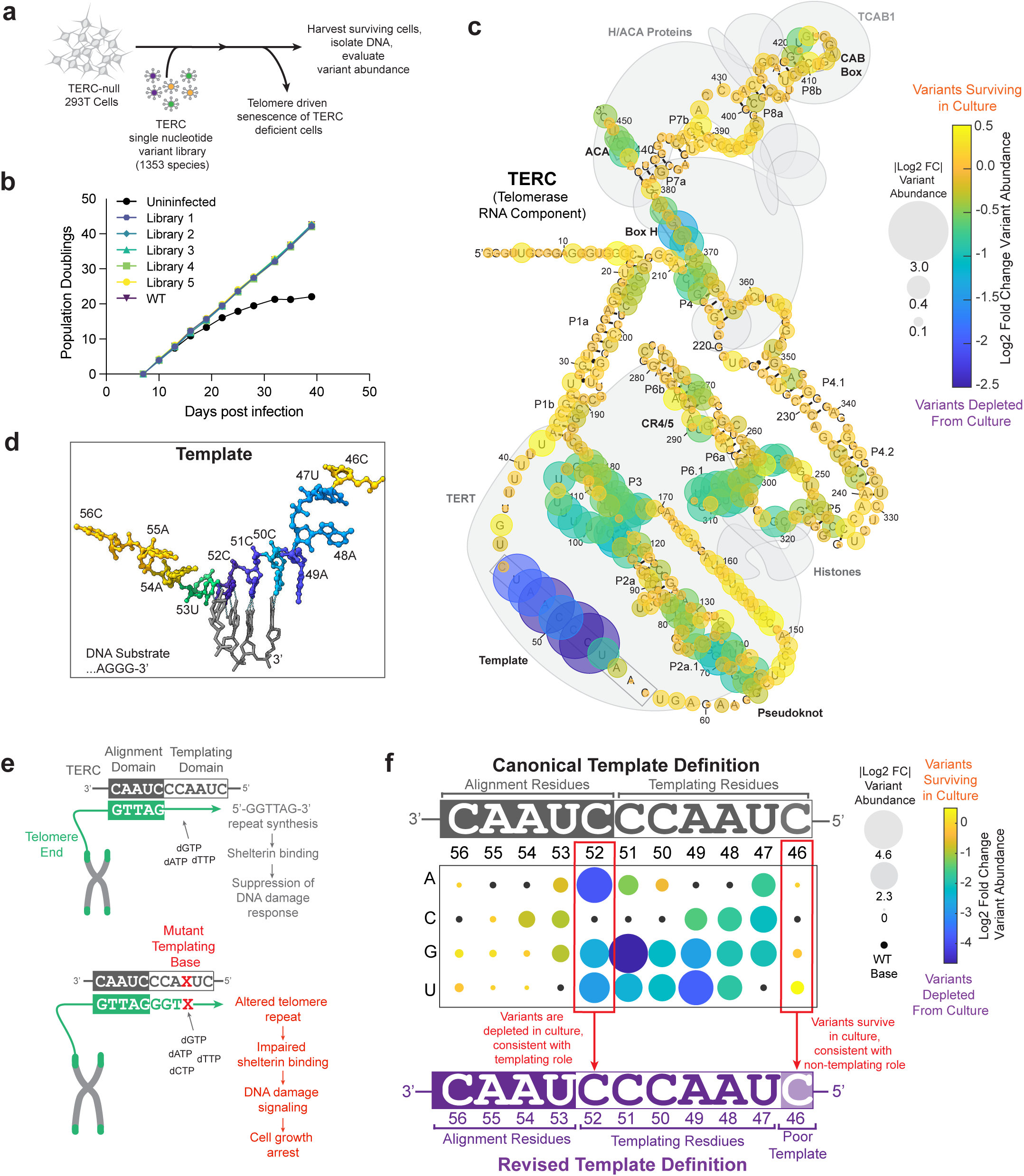
Saturating variant analysis of the human telomerase RNA component. **a,** Diagram of TERC variant screening strategy. **b,** Growth curve of 293T TERC-null cells infected with the indicated lentiviral TERC expression vector compared with uninfected cells. Representative data shown from one of two biological replicates. **c,** Display of TERC saturating mutagenesis results. Mean log_2_[fold change] in variant abundance is calculated by comparing variant abundance in TERC-null 293T cells after 40 days of culture versus variant abundance in plasmid libraries. Variant effects are averaged for a given base and overlaid onto a schematized structure of TERC (adapted from ref. ^32^). Color indicates log_2_[fold change]. Dot size indicates absolute value of log_2_[fold change]. Data represent the average of two biological replicates. **d,** Data from **c** overlaid onto the Cryo-EM structure of the TERC template and DNA substrate within the telomerase active site. Structure from PDB 9qax^32^. Color scale is the same as used in **c**. **e,** Schematic depicting how TERC templating base mutations have deleterious effects. **f,** Display of TERC saturating variant analysis from **c** demonstrating the relative dropout of single nucleotide variants across the template domain. Color indicates log_2_[fold change]. Dot size indicates absolute value of log_2_[fold change]. Black dots indicate wildtype bases. Data represent the average of two biological replicates.

### Discordant effects of annotated TERC template and alignment domain variants

The variants most depleted in our screen mapped to the template domain, comprised of the canonically annotated alignment and templating residues (**Fig. 1c**, Extended Data Fig. 2a,e,g). Mutation of TERC templating residues can introduce alternative sequences into telomeres that disrupt the shelterin complex and lead to DNA damage signaling, senescence-associated changes, and cell growth arrest in a dominant fashion, independent of telomere length (**Fig. 1e**)^34,35^. The canonical annotation of the human telomerase RNA template defines the templating residues as n.46-51 and alignment bases as n.52-56 (**Fig. 1f**). We were thus surprised to find that variants at position 46 were well tolerated in the screen, while position 52 mutations were the most negatively selected of all 451 positions in the TERC gene (**Fig. 1f**, Extended Data Fig. 2g). These data led us to hypothesize that bona fide TERC templating function in human cells may be encoded at positions n.47-52 rather than the long-standing annotation n.46-51 (**Fig. 1f**). To rigorously interrogate the templating function of TERC positions 46 versus 52 (**Fig. 2a**), we generated lentiviral expression vectors encoding all single nucleotide variants at these positions and tested their effects in telomerase-deficient cells. In addition to growth arrest (**Fig. 1a**), TERC-null 293T cells show characteristic morphologic signs of senescence with critical telomere attrition or dysfunction, as well as phosphorylation of CHK2 (pCHK2) as an indicator of the DNA damage response^30^. We thus evaluated effects of overexpressing TERC variants at positions 46 vs 52 in TERC-null 293T cells with respect to: 1) cell growth, 2) senescence-associated morphological changes, and 3) DNA damage signaling (**Fig. 2b**). We found that the introduction of vectors encoding TERC position 52 variants strongly impaired cell growth compared to uninfected cells (**Fig. 2c**). By brightfield microscopy, position 52 TERC variants also rapidly drove morphologic hallmarks of senescence including syncytia formation (**Fig. 2d**, Extended Data Fig. 3a) well before the onset of these changes in uninfected TERC-null cells. In contrast, position 46 variants rescued TERC-null 293T cells from short telomere-induced growth arrest indistinguishably from WT TERC (**Fig. 2b**) and did not cause morphologic signs of senescence (Extended Data Fig. 3a). These data suggested that position 52 variants have a dominant negative effect from templating the wrong base, whereas 46 variants show no alterations or impairments in the ability to synthesize WT telomere repeats. Next, to assess for DNA damage signaling in TERC-null cells expressing TERC position 46 or 52 variants, we performed immunoblotting for pCHK2 and found that position 46 variants reduced pCHK2 signaling in near-senescent TERC-null 293T cells in a manner similar to WT TERC, again consistent with position 46 variants having the capacity to synthesize and restore WT telomere repeats (**Fig. 2e**). In contrast, position 52 mutants failed to reduce pCHK2 signaling. Collectively, in showing that position 52 TERC variants are cytotoxic while position 46 variants act similarly to WT TERC and efficiently rescue TERC-null 293T cells from senescence, these data validate the screening results and point to a functional reannotation of the TERC template domain.

**Fig. 2.**
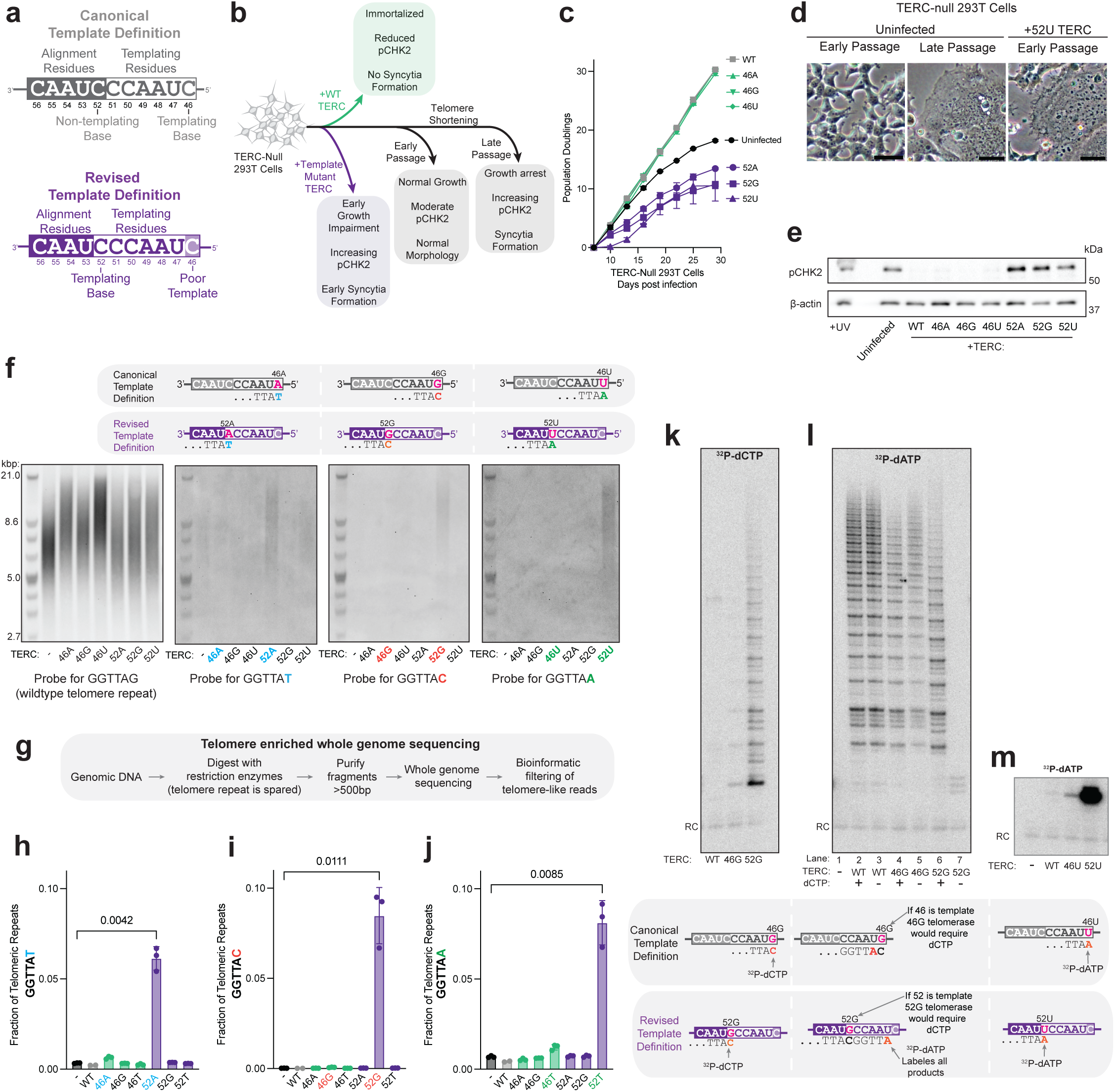
Cellular and enzymatic evidence for a redefined telomerase template. **a,** Schematic depicting canonical and revised template definitions. **b,** Diagram depicting the effects of wildtype versus template mutant TERC expression in TERC-null 293T cells. **c,** Population doubling curves of uninfected TERC-null 293T cells compared with cells infected with the indicated lentiviral TERC vectors. Two biological replicates are displayed. Error bars indicate range. **d,** Representative brightfield microscopy images of early passage and late passage TERC-null 293T cells compared with 52U TERC expressing TERC-null 293T cells from **c**. Early passage images acquired 15 days post infection. Scale bar 50µm. **e,** Immunoblot of cells from **c**, probing for pCHK2, and beta actin. UV irradiated 293T cells were used as a positive control. Cells were 19 days post infection. One representative blot is shown from three biological replicates. **f,** TRF Southern blots of gDNA from 293T cells 19 days after infection with the indicated TERC lentiviral vectors. The same blot was stripped and probed sequentially for the indicated targets. One representative set of blots is shown from three biological replicates. **g,** Strategy for telomere enriched whole genome sequencing. **h-j,** Quantification of the fraction of the indicate telomere variant repeats out of the total telomeric repeats in telomeric reads performed by processing gDNA from **f** as described in **g** (see methods). Experiment performed in biological triplicate. Error bars indicate SD. P value calculated using paired two-sided T-test. **k-m,** Direct telomerase assay performed using telomerase immunopurified from TERC-null 293T cells transfected with vectors encoding tagged TERT and the indicated TERC variant. Reactions include immunopurified telomerase, 18-TTA primer and ^32^P-dATP or ^32^P-dCTP as indicated. In **k** and **m**, 5 µM each of dCTP, dATP, dGTP, and dTTP was used. In **l**, 5 µM each of dATP, dGTP, and dTTP was used, with addition of 5 µM dCTP as indicated. RC = radiolabeled 15 nt recovery control oligo.

## Genomic evidence for a redefined telomerase RNA template domain

We next sought to directly detect variant sequences in native telomere ends after expression of position 46 versus 52 variants. Based on the results above, we hypothesized that position 46 variants would be able to elongate telomeres with a sequence composition and extent similar to WT, whereas position 52 variants would introduce variant sequences with consequent defects in telomere lengthening. We thus stably introduced lentiviruses encoding TERC position 46 and 52 variants into 293T cells and assessed for changes in telomere length by Southern blotting using a probe to detect WT repeats. We found a clear increase in mean telomere length from co-expression of position 46 variants with endogenous TERC, whereas position 52 variants appeared to expand the heterogeneity of telomere length without increasing mean telomere length (**Fig. 2f**, Extended Data Fig. 3b). When we used variant-specific Southern blot probes, we found that position 52 TERC variants were indeed creating altered repeat sequences on native telomere ends, in contrast to position 46 variants which showed no hybridization signals corresponding to variants (**Fig. 2f**). Next, we performed telomere-enriched whole genome sequencing on 293T cells expressing TERC position 46 or 52 variants to precisely quantify variant repeat incorporation (**Fig. 2g**). Filtering for stretches of telomere-like reads and quantifying wildtype and variant telomeric repeats, we found that expression of position 52 variants led to strong increases in the abundance of the corresponding variant telomere repeats whereas position 46 variants did not (**Fig. 2h-j**), consistent with the Southern blotting results. Collectively, these data conclusively demonstrate that position 52 of TERC serves a core templating function for telomerase in human cells, whereas position 46 is dispensable.

### In vitro evidence for redefinition of telomerase RNA templating residues

In vitro studies of human telomerase over the past 30 years have yielded significant insights into its reverse transcription mechanisms including functional interrogation of the TERC template domain, but have not systematically questioned templating residue annotation. We thus sought to reconcile and extend our studies indicating a reassignment of the TERC templating residues using in vitro telomerase assays. In the direct telomerase assay, reconstituted or purified telomerase is incubated with primers representing telomeric repeats, plus radiolabeled and cold deoxynucleotides, followed by direct visualization of telomerase extension products by denaturing polyacrylamide gel electrophoresis. To generate purified telomerase harboring variant templates for these in vitro studies, we transfected TERC-null 293T cells with a vector encoding FLAG-TERT plus a vector encoding WT TERC or TERC position 46 or 52 variants. Telomerase was immunopurified from cells using anti-FLAG beads followed by elution with FLAG peptide. Using the direct telomerase assay, we first compared wildtype TERC, 52C>G TERC (52G), and 46C>G TERC (46G) for their ability to incorporate ^32^P-dCTP. In this experimental setup, the relative efficiency of templating activity at position 52 and 46 is reflected in the extension of telomeric primers by ^32^P-dCTP, which will only be incorporated across from the variant G residue. We found that WT telomerase showed no radiolabeled repeat synthesis as expected, as the WT template does not contain a templating G residue and does not use deoxycytidine nucleotides (**Fig. 2k**). In contrast, we found that 52G telomerase robustly synthesized ^32^P-dCTP containing repeats compared to 46G telomerase (**Fig. 2k**). Next, we evaluated 52G and 46G TERC in the direct telomerase assay using ^32^P-dATP plus cold dATP, dGTP, and dTTP, with or without dCTP. Using all dNTPs, we found that 46G and 52G variant telomerases efficiently produced extension products of a similar pattern but reduced processivity compared with wildtype TERC (**Fig. 2l**, lanes 2,4,6). Removing dCTP from the reaction had minimal effect on the overall activity of wildtype TERC or 46G TERC, whereas the absence of dCTP rendered 52G TERC completely inactive (**Fig. 2l**, lanes 3,5,7). This finding indicates that the majority of 46G extension products do not contain dCTP, clearly demonstrating that position 46 is not used as a core templating residue in vitro. In contrast, all extension products of 52G TERC depended on the presence of dCTP, indicating that position 52 is the dominant templating residue. Finally, we compared wildtype TERC and TERC variants 46C>U (46U), and 52C>U (52U) using a primer substrate ending in TTA-3’ and ^32^P-dATP. In this experimental setup, WT telomerase incorporates a cold dGTP residue as the first base, while 46U or 52U variant TERCs could incorporate radiolabeled dATP as the first templated nucleotide depending on whether the variant residue was used as a templating base. We found that ^32^P-dATP was incorporated as the first base by 52U TERC >100 times more efficiently compared with 46U TERC (**Fig. 2m**). Collectively, these data both in vitro and in cells rigorously demonstrate that position 52 is a dominant templating base responsible for telomere repeat synthesis by human telomerase, a function that was previously attributed to position 46.

## Flexible templating residue usage by human telomerase

In our in vitro studies, we found that although position 52 is the primary templating base for telomerase, position 46 is not entirely devoid of templating activity (**Fig. 2k, m**). This observation raised the question of whether position 46 is ever used as a templating base by telomerase in human cells, which would deviate from the existing model in which only hexamers are created. To address this, we first asked if position 46 template usage could be detected in our telomere-enriched whole genome sequencing datasets of genomic DNA from 293T cells expressing position 46 and 52 TERC variants (**Fig. 2g**). We hypothesized that position 46 templated variant repeat sequences, if they existed at all, would occur in isolation flanked by WT telomeric sequences, given the low templating frequency of position 46 observed in the direct telomerase assay. We thus adjusted our NGS analysis filter and quantified the occurrence of single variant repeats flanked by WT repeats in position 46 variant TERC-expressing cells (**Fig. 3a,b**). Using these parameters, we now detected a significantly increased representation of single variant repeats corresponding to the specific position 46 variant introduced into the cells (**Fig. 3c-e**). Further analysis of the sequence context of position 46 and 52 encoded variants confirmed that position 46 variants were most likely to occur as single variants surrounded by wildtype telomere repeats, whereas position 52 encoded variants were very likely to be followed by another variant repeat, commonly occurring in tandem runs of 8 or more (Extended Data Fig. 4a). These data reveal for the first time that human telomerase has flexible template usage beyond a hexamer, with positions 52-47 serving as dominant templating bases but also including position 46 as a low efficiency templating base, both in vitro and in cells.

**Fig. 3.**
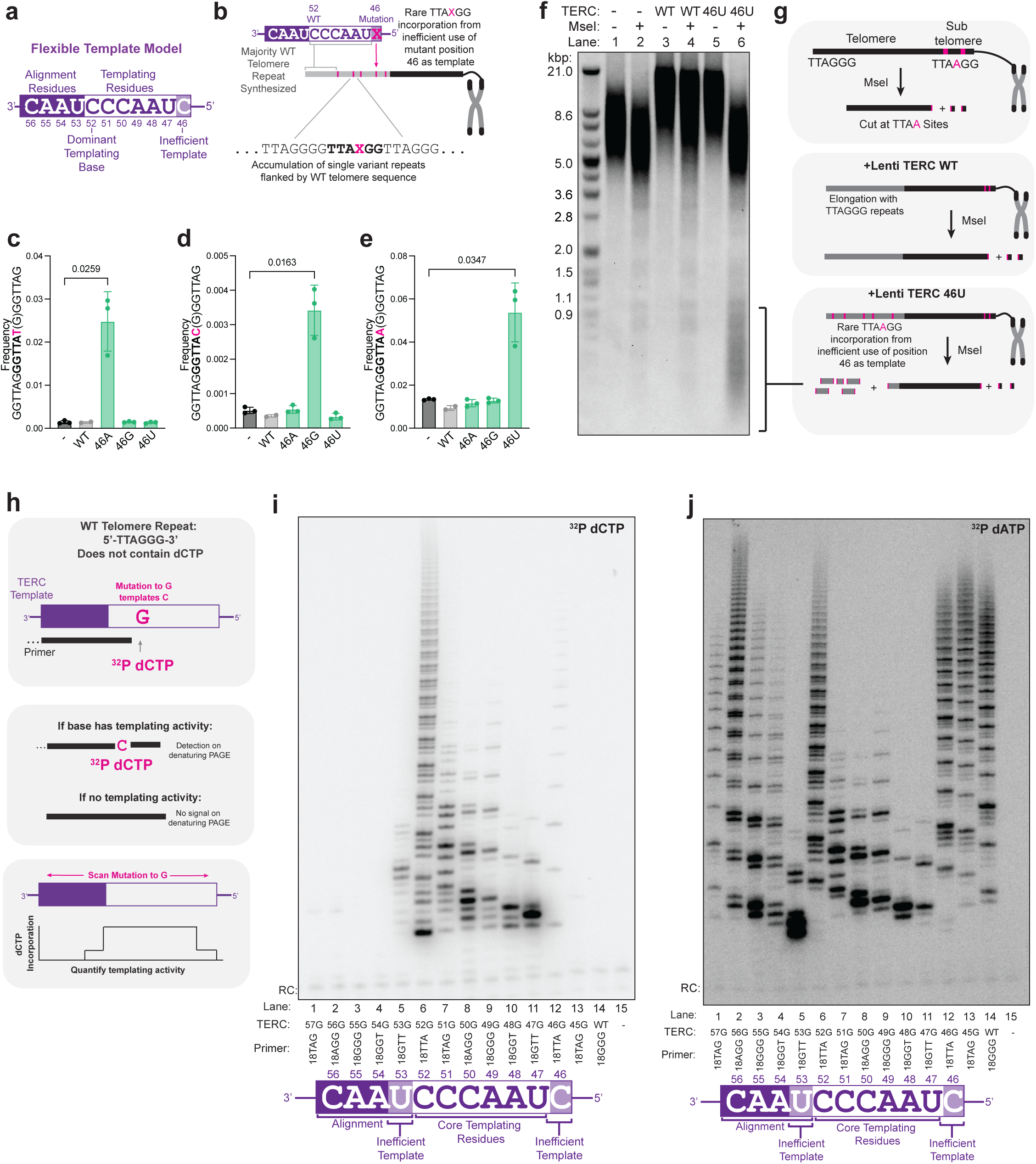
Flexible use of eight templating bases by human telomerase. **a,** Schematic of TERC template domain with proposed domain definitions. **b,** Schematic of how flexible template use of position 46 can lead to incorporation of single mutant telomeric repeats flanked by wildtype repeats. **c-e,** Analysis of telomere enriched whole genome sequencing from Fig. 2h-j, quantifying the frequency of the indicated singleton telomere variant repeat present in telomere-like reads per million total reads. P value calculated using paired two-sided T test. Error bars indicate SD. **f,** TRF Southern blot of DNA from 293T cells infected with the indicated lentiviral TERC constructs and cultured for 19 days, performed with or without MseI as indicated. One representative blot is shown from two biological replicates. **g,** Diagram of how MseI digest allows for detection of rare 46U TERC encoded variants. **h,** Diagram of G-scan mutagenesis strategy to identify TERC templating residues. **i,** Direct telomerase assay performed using telomerase immunopurified from TERC-null 293T cells transfected with vectors encoding tagged TERT and the indicated TERC variant. Reactions include immunopurified telomerase, 5 µM each of dCTP, dATP, dGTP, and dTTP, the indicated primer, as well as ^32^P-dCTP. RC = radiolabeled 15 nt recovery control oligo. **j**, Direct telomerase assay performed as in **i** using ^32^P-dATP.

The lack of detection on Southern blot of position 46 variant-encoded repeats (**Fig. 2f**) can be explained both by their rarity and also because tandem repeats are required for efficient hybridization of variant-specific probes. Thus, to directly visualize the flexible use of TERC position 46 as a templating base in human cells, we leveraged the fact that the variant repeat potentially encoded by 46U, 5’-GGGTTA<u>A</u>-3’, would contain a 5’-TTAA-3’ sequence that is not otherwise present in the telomere, and which is susceptible to cleavage by the MseI restriction enzyme. We hypothesized that restriction digest with MseI would lead to detectable trimming of telomere length on Southern blot in TERC 46U expressing cells due to rare incorporation of 5’-GGGTTA<u>A</u>-3’ repeats in newly extended telomeres. We thus performed Southern blotting on genomic DNA from 293T cells stably expressing lentiviral WT or 46U TERC for 19 days, followed by probing for the WT telomere repeat (**Fig. 3f,g**). We found elongation of telomeres in cells expressing 46U or WT TERC, consistent with our prior results (**Fig. 3f**, lanes 1,3,5). However, when we added MseI to the restriction digest, we found a reduction of the extended telomeres from TERC 46U expressing cells back to baseline (**Fig. 3f**, lanes 6 versus 2), accompanied by the appearance of a robust telomeric signal of mean average length < 1kb. These low molecular weight fragments represent stretches of WT telomere repeats flanked by rare 46U encoded repeats that have been cleaved off the telomere end (**Fig. 3g**). We noted the size of the low MW fragments as several hundred nucleotides long (or ∼100x telomeric repeats that are 6 nt long), yielding an estimate that 46U is used as a templating residue approximately ∼1/100^th^ of the time, in line with in vitro results (**Fig. 2m**). Similar results were found in K562 cells (Extended Data Fig. 4b). Collectively, these data conclusively demonstrate that human telomerase shows flexibility in the templating function of TERC beyond hexamers, with position 46 infrequently utilized as a templating residue in human cells.

## The range of templating bases used by human telomerase

Based on our findings that position 52 is an undescribed templating base and that human telomerase shows flexible templating, we wondered whether additional TERC residues could act as templating bases. To address this, we generated TERC expression vectors harboring point mutations across template domain residues n.45-57, changing each base individually to G (hereafter termed “G-scan variants”). As deoxycytidine is not present in the wildtype telomere repeat, the use of dCTP by G-scan variants reflects the templating activity of only the given mutant base (**Fig. 3h**). We expressed G-scan variants in TERC-null 293T cells with FLAG-TERT and immunopurified the variant telomerases. We then evaluated the telomerase activity of these variants in the presence of all four cold dNTPs plus ^32^P-dCTP, and telomere repeat primers that hybridize at the base immediately before the G-scan variant (**Fig. 3i**). Remarkably, we found that position 53, previously annotated as an alignment base, could also serve as a templating base for telomerase in vitro. As a control to avoid false negative interpretation of the lack of dCTP incorporation, we utilized ^32^P-dATP instead of ^32^P-dCTP and verified the activity of all immunopurified telomerase variants (**Fig. 3j**). All in all, G-scan mutants at positions n.53-46 had detectable incorporation of dCTP (**Fig. 3i**, lanes 5-12), with position n.52-47 mutants being the most active for incorporating dCTP (**Fig. 3i**, lanes 6-11). These data align with our screening results indicating that variants at positions n.52-47 are the most toxic to TERC-null cells (**Fig. 1c**). Of note, position 53 variants were more negatively selected in our screen than position 46 variants, although less negatively selected than mutations to positions n.52-47 (**Fig. 1f**). Together, these data support the conclusion that TERC positions n.52-47 comprise the core template, but also unexpectedly show that human telomerase can use a range of its associated RNA residues including n.53 and/or n.46 for flexible templating.

With these insights in hand, we again turned to whole genome sequencing data to search for instances of flexible telomerase template usage in human cells. We reasoned that occasional usage of an altered templating base at position 46 would mark the end of one catalytic cycle and allow for detection of position 53 templating activity in the subsequent catalytic cycle. The most likely possibility of detecting this would be if base-pairing was still possible at n.54A, and thus if the variant at position 46 was also A (Extended Data Fig. 4c). Indeed, we found that after the incorporation of an n.46A encoded variant T at the telomere end, the subsequent sequence was occasionally 5’-<u>A</u>GGGTTA-3’, indicating that position 53 was used as a templating base to encode the 5’-most A residue in that sequence (Extended Data Fig. 4c-e). Remarkably, in the case of 46A TERC, we reproducibly observed a rare instance of the sequence 5’-AGGGTTAT-3’, which would be encoded if both residues 53 and 46 were utilized in a single primer extension event (Extended Data Fig. 4c-d,f). Collectively, these findings indicate that telomerase can flexibly use not six but eight nucleotides in the TERC template domain as bona fide templating bases in human cells.

## Identification of hyperactive TERC variants with an abbreviated alignment domain

During G-scan mutagenesis in the ^32^P-dATP controls, we unexpectedly found that mutation of position 56 dramatically increased maximal telomerase product size compared to WT TERC and other variants (**Fig. 3j**, lane 2). Position 56 is canonically the 3’ most nucleotide of the alignment domain, which re-binds the telomere end after each round of repeat synthesis. We were thus surprised to find increased activity in the 56G variant, as mutations in the alignment domain have previously been shown to impair telomerase processivity^36^. To further characterize how mutation of position 56 impacts telomerase activity, we generated TERC expression vectors encoding 56A and 56U and tested their activities alongside 56G and wildtype TERC (encoding 56C) in the direct telomerase assay. We found that the products formed by any position 56 variant had a greater maximum size than WT TERC at each assay timepoint (**Fig. 4a**). We next asked whether position 56 variants showed increased telomerase activity compared to WT TERC in human cells. Indeed, we found that all position 56 variants reproducibly led to significantly greater telomere lengthening than WT TERC when overexpressed in 293T TERC-null cells (**Fig. 4b-c**). Given that any position 56 mutation resulted in increased activity, we asked how deletion of n.56 (56Δ) affected telomerase activity in cells. We found that 56Δ also increased telomere length when compared with WT TERC transiently over-expressed in cells (**Fig. 4d-e**). We next asked whether position 56 variants could drive enhanced telomere lengthening in cells derived from patients with telomere biology disorders. Overexpressing position 56 variant TERC in cells from TBD patients harboring two different mutations in *DKC1*, a telomerase complex protein essential for TERC stability, again led to strong increases in telomere length beyond the effect from wildtype TERC expression (**Fig. 4f-h**). Together, these data indicate that an abbreviated alignment domain with mutation or deletion of position 56 enhances human telomerase activity, thus revealing the first hyperactive human TERC variants.

**Fig. 4.**
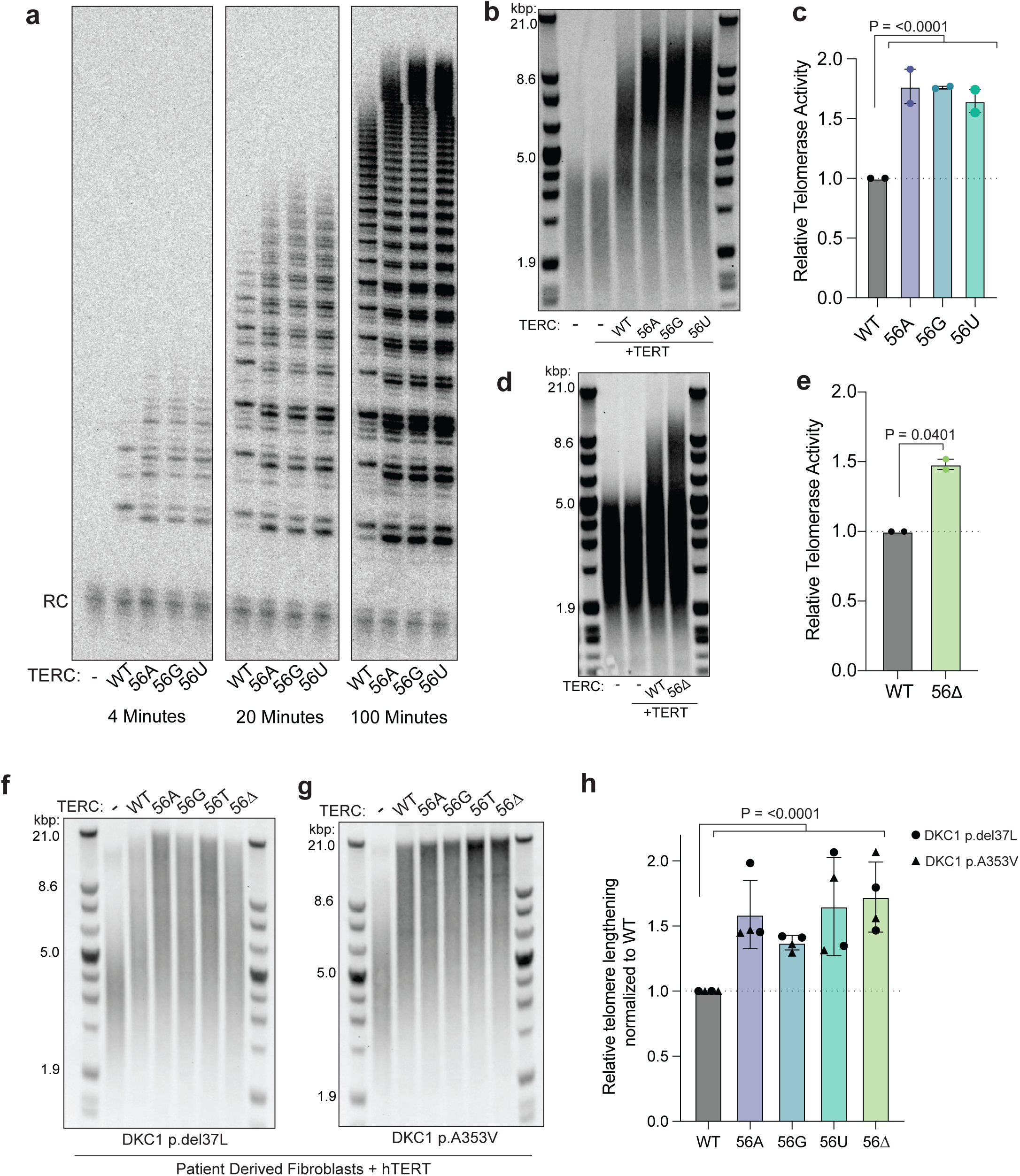
Identification of hyperactive TERC variants with an abbreviated alignment domain. **a,** Direct telomerase assay performed using telomerase immunopurified from TERC-null 293T cells transfected with vectors encoding tagged TERT and the indicated TERC variant. Reactions include immunopurified telomerase, 5 µM each of dCTP, dATP, dGTP, and dTTP, 18-GGG primer, as well as ^32^P-dATP. Reactions performed for the indicated time period. RC = radiolabeled 15 nt recovery control oligo. **b,** TRF Southern blot of TERC-null 293T cells transfected with pBS-U3-TERC and TERT vectors as indicated followed by culture for 48 hrs. A representative blot is shown from two biological replicates. **c,** Quantification of **b**. Error bars indicate range. P value calculated using paired two sided test. **d,** TRF Southern blot of TERC-null 293T cells transfected with pLKO.1-U3-TERC and TERT vectors as indicated followed by culture for 48 hrs. A representative blot is shown from two biological replicates. **e,** Quantification of **d**. **f, g** TRF Southern blot of the indicated cell lines infected with the indicated lentiviral TERC expression vector. Two biological replicates were performed: one cultured for two weeks (displayed), and one for four weeks. **h,** Quantification of two biological replicates from **f** and **g**. Error bars indicate range. P values in this figure were calculated by paired two sided t test performed testing if log2 fold change values for variants are greater than 0 (relative telomere lengthening greater than 1).

## Mammalian evolution of an expanded template domain and altered template flexibility

Position 46 of TERC is one of the most highly conserved template domain residues across vertebrates (**Fig. 5a**, orange highlight)^33^. Why would position 46 be conserved in the template domain if it plays such a negligible role as a templating residue in human telomerase? Along the same lines, why would an alignment base at position 56 constrain telomerase activity? Upon phylogenetic comparison, we found that the 4 nt telomere sequence alignment region in most vertebrates like fish and reptiles (**Fig. 5a**, green) appears to have undergone an expansion to 5 nt with the introduction of cytosine corresponding to position 56 in human telomerase RNA (**Fig. 5a**, blue) and certain other mammals. We wondered if this alignment domain expansion may have impacted the utilization of position 46 as a templating base (**Fig. 5b**). This model could explain why position 46 is so highly conserved despite being relatively unimportant for human telomerase activity – position 46 transitioned from a dominant templating base to an infrequently used templating residue because of alignment region expansion relatively recently in mammalian evolution.

**Fig. 5.**
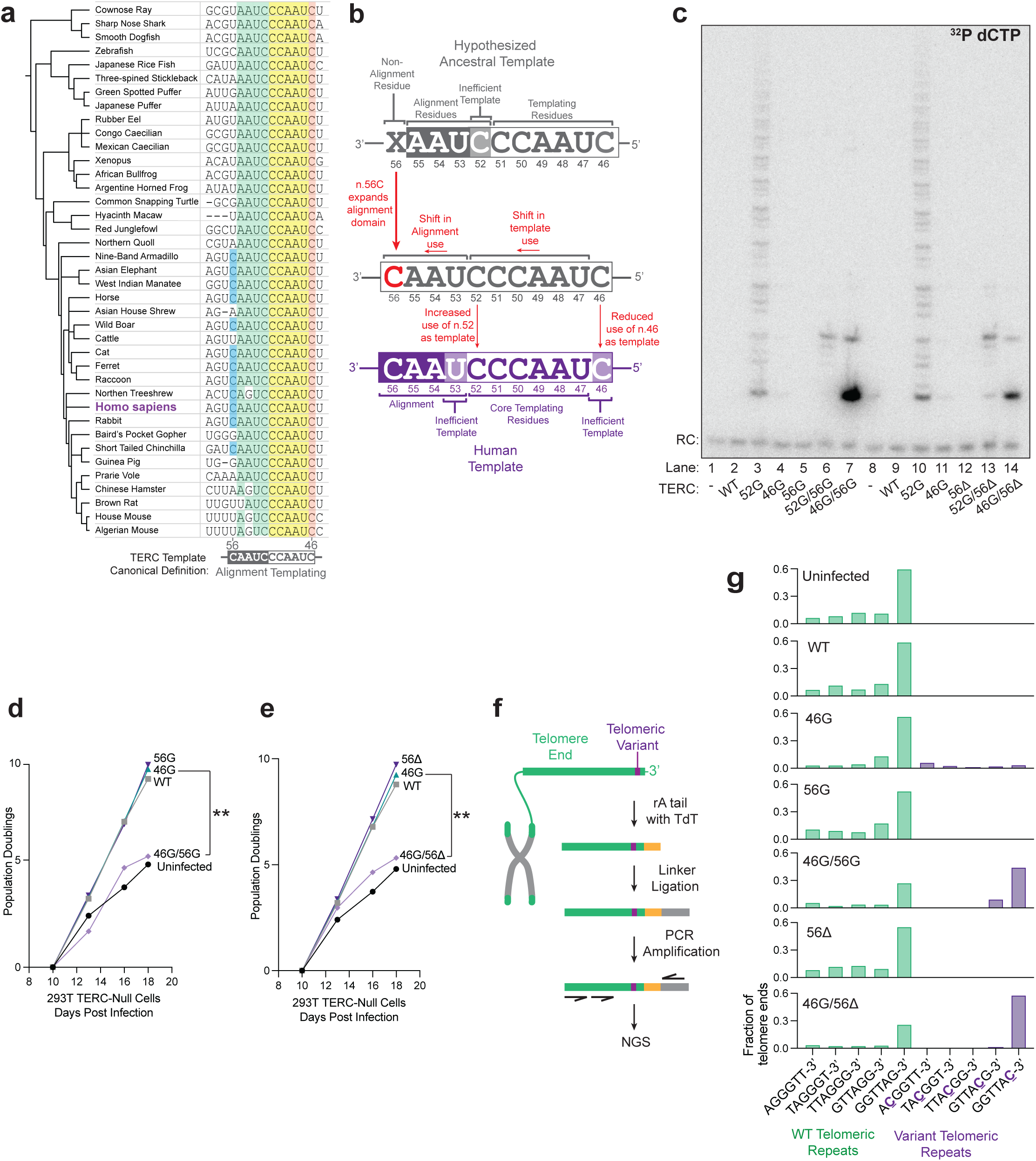
A template expansion in mammalian evolution shifted template usage. **a,** Vertebrate phylogenetic tree of TERC template domains. Numbering and alignment relative to human TERC. n.56C is highlighted in blue. Other canonical alignment region bases n.55-52 are highlighted in green. Canonical templating bases n.51-47 are highlighted in yellow. Position 46 is highlighted in orange. **b,** Diagram describing how mutation of position n.56C could lead to a shift in templating register. **c,** Direct telomerase assay performed using telomerase immunopurified from TERC-null 293T cells transfected with vectors encoding tagged TERT and the indicated TERC variant. Reactions include immunopurified telomerase, 5 µM each of dCTP, dATP, dGTP, and dTTP, 18-GGG primer, as well as ^32^P-dCTP. RC = radiolabeled 15 nt recovery control oligo. **d-e,** Population doubling curves of uninfected TERC-null 293T cells compared with cells infected with the indicated lentiviral TERC expression vectors. P value for difference in slopes calculated by linear regression. **f,** Diagram of telomere end sequencing approach (see Methods). **g**, Fraction of telomere ends terminating in the indicated 6-mer repeat determined by telomere end sequencing of TERC-null 293T cells expressing the indicated lentiviral TERC expression vector for 9 days.

To test this hypothesis, we used multiple approaches to examine the impact of position 56C on the utilization of position 46 as a templating base. First, we interrogated the effects of variations at TERC position 56 on the ability of TERC variant 46G to drive templating of ^32^P-dCTP in the direct telomerase assay. We found that in contrast to the poor dCTP utilization by TERC 46G alone (**Fig. 5c**, lanes 4 and 11), the double mutants TERC 46G/56G and TERC 46G/56τι showed strong dCTP incorporating activity in the products (**Fig. 5c**, lanes 7 and 14), indicating markedly increased usage of position 46 as a templating residue. In keeping with a shift of template residues rather than expansion of the templating domain, we found that whereas TERC 52G alone showed strong ^32^P-dCTP utilization and processivity, this activity was abrogated by simultaneous variation at position 56 (i.e., 52G/56G and 52G/56Δ; **Fig.5c**, lanes 6 and 13 compared to lanes 3 and 10)). These data are consistent with the hypothesis that the additional capacity for aligning the telomere end at position 56C shifted TERC templating function from predominantly using position 46 to using position 52. To examine these effects in human cells, TERC-null 293T cells were infected with lentiviral vectors encoding TERC 46G versus TERC 46G plus position 56 mutations. We hypothesized that if position 46 gained templating activity by simultaneous disruption of position 56, position 46 variants would now compromise cell growth due to aberrant variant repeat incorporation into the telomere, instead of rescuing cell growth by synthesizing predominantly WT telomere repeats as they did before (**Fig. 2c**). In these experiments, we found that TERC variant 46G alone efficiently rescued cells from senescence in a manner similar to WT TERC (**Fig. 4i-j**), as expected from our previous results (**Fig. 2c**). TERC variants 56G or 56τι alone were also able to rescue cells from senescence (**Fig. 5d-e**), consistent with their intact capacity for telomere elongation with WT telomeric repeats (**Fig. 4b-e**). However, we found that when these residues were simultaneously disrupted, neither position 46G/56G nor position 46G/56τι TERC variants could rescue TERC-null 293T cells from senescence (**Fig. 5d-e**). These results are consistent with altered telomerase activity without restoration of functional telomeres. We predicted this altered telomerase activity would be due to increased use of position 46G as a templating residue that would disrupt telomere repeat addition at the chromosome end by incorporation of a terminal C residue. To investigate this, we amplified the telomere ends from genomic DNA after linker ligation and performed NGS to characterize the terminal 6-mer sequence (**Fig. 5f**). Strikingly, and consistent with our hypothesis, we found that a majority of telomere ends in TERC-null 293T cells expressing 46G/56G and 46G/56τι TERC variants now encoded a terminal cytosine residue (**Fig. 5g**). Together, these data indicate that genetic variation in the alignment domain of TERC influences core templating residue function as well as flexible template usage in human telomerase. The results are consistent with the hypothesis that an early mammalian ancestor of humans underwent an expansion of the alignment domain that constrained telomerase reverse transcriptase activity through base pairing in the alignment region, and shifted position 46 templating activity to position 52.

## Discussion

Telomerase plays critical roles in cell function and human health. TERC emerged as one of the first long non-coding RNAs associated with disease, amplified in cancers and mutated in Mendelian TBDs^18,37–39^. Intensive studies of TERC structure and biogenesis have yielded insights into its essential functions as a template and scaffold of telomerase, informing our basic understanding of this unique DNA polymerase and opening diagnostic and therapeutic avenues. Here, we report saturating variant analysis of TERC, the first to our knowledge for a long ncRNA, and using cell-based functional readouts show its template has been misannotated in the 30 years since its cloning. We relocate the canonical telomere repeat templating core and reveal that human telomerase flexibly uses a range of nucleotides for reverse transcription. Integrating these findings, we propose a new mechanistic model for telomerase reverse transcriptase activity, a critical enzymatic process for maintaining genome integrity at the ends of all human chromosomes (**Fig. 6**).

**Fig. 6.**
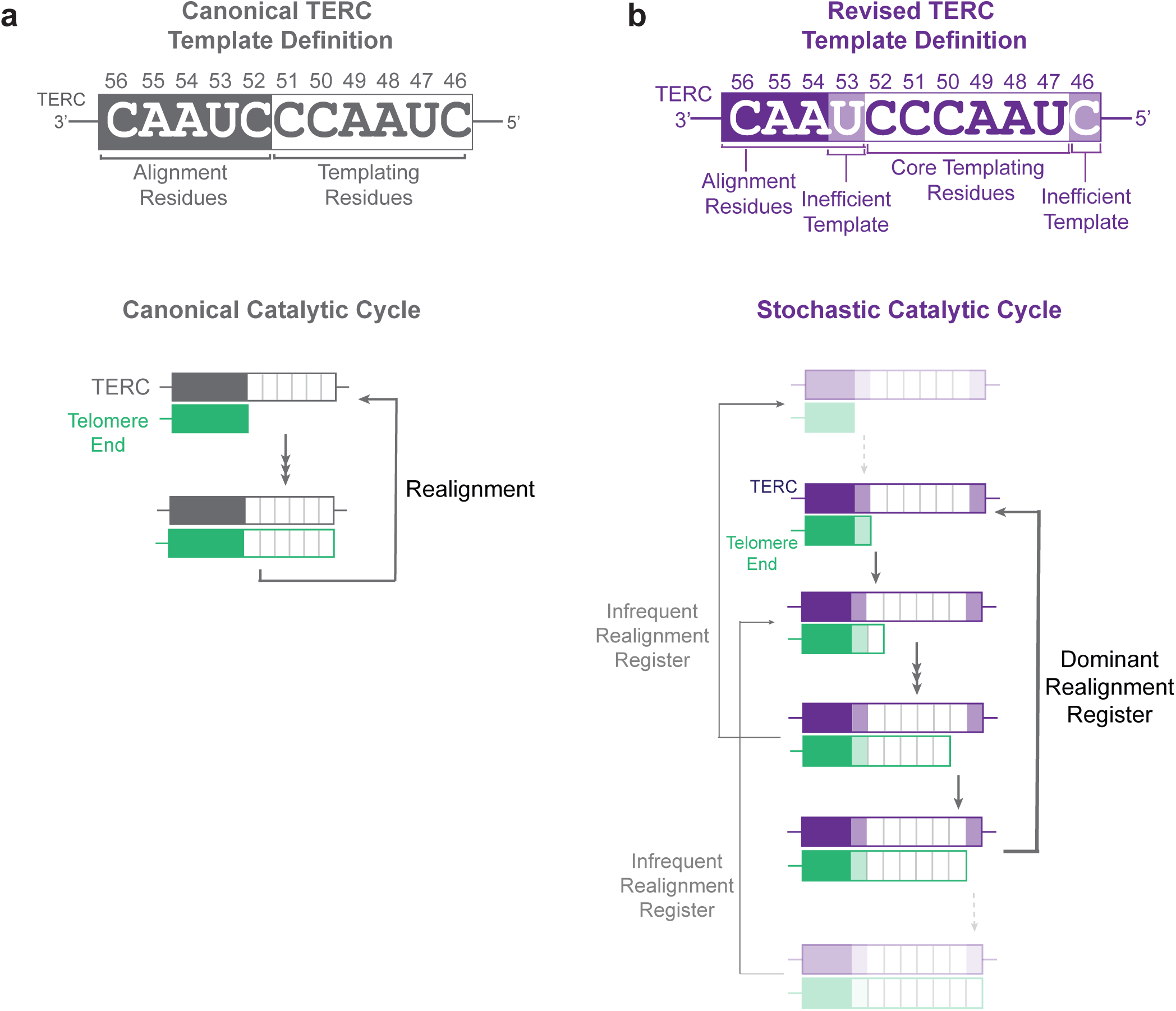
A revised, flexible model for the telomerase catalytic cycle. **a,** Canonical model for the telomerase catalytic cycle. Telomerase extends the telomere end by six nucleotides templated by positions n.51-46. After repeat synthesis, the newly extended telomere end realigns to base pair with the alignment domain (positions n.56-52 of TERC), enabling processive repeat synthesis. **b,** Revised model for the telomerase catalytic cycle. During repeat synthesis, the telomere end can be extended by reverse transcription templated by TERC positions n.53-46, with positions n.52-47 being the predominant templating bases. Within a given round of repeat synthesis, as the end of the template is approached there is an increasing probability of realignment, with the most common realignment register being after synthesis templated by position n.47 when the telomere ends in TTA-3’ (bold arrow). Infrequently, realignment can occur one position earlier or later (faint arrows). If realignment occurs early when the telomere ends with GTT-3’, position n.53 can be used as a templating base on the subsequent catalytic cycle. Realignment can also occur late, following position n.46 use as a template. Within this model, between 4-8nt can be synthesized in a given primer extension event, with 5’-GGGTTA-3’ being the most common extension event.

The data presented revise several fundamental tenets of the prevailing model of the telomerase template domain and catalytic cycle (**Fig. 6**). To date, telomere repeat synthesis by human telomerase has been defined as 1) incorporating precisely **six** nucleotides per catalytic cycle, 2) in the order 5’-**GGTTAG**-3’, 3) templated by **n.46-51** of TERC, 4) followed by dissociation and re-annealing to a **5 nt** alignment sequence n.52-56, 5) the extent of which is **critical for its activity** and processivity. The model emerging from our data (**Fig. 6B**) argues instead that 1) up to **eight** nucleotides can be flexibly templated per catalytic cycle, 2) with 5’-**GGGTTA**-3’ as the dominant six nucleotide repeat, 3) templated by **n.47-52** of TERC, 4) followed by translocation to a **4 nt** alignment domain n.53-56, 5) with base pairing at position 56 **constraining telomerase activity**. In light of our findings, what explains the origin of the long-held mechanistic model of human telomerase templating function? We suggest that inferences from in vitro ciliate telomerase studies and phylogenetic conservation may have skewed the early functional annotation of the TERC template domain. In 1989, upon detecting telomerase activity from human cells for the first time, Morin sequenced the 6 nt ladder products and found they predominantly ended in GGTTA**G**-3’^21^. Two possible explanations were raised: 1) there was a catalytic barrier between incorporation of the first and second of the three consecutive G residues in each repeat synthesis cycle, or 2) the templating boundary of human telomerase RNA would end at a 5’-C in the sequence 5’-CUAACC-3’, requiring translocation of the elongating telomere end and/or its release, thus resulting in accumulation of the observed sequence. The second scenario was favored based on logical inference, as well as *Tetrahymena* studies where telomerase RNA templating boundary had been characterized^40^. The subsequent cloning of human TERC and other vertebrate telomerase RNAs indeed revealed a conserved 5’-C in this context (position 46 in humans), reinforcing the prevailing model wherein the 6 templating residues of TERC are located at positions 51-46^6,22^. However, in the decades since this model of TERC templating function was proposed, to our knowledge it was not rigorously put to the test in a cellular context.

Here, prompted by the results of unbiased screening in human cells, we comprehensively re-evaluated the TERC template domain and relocated its dominant templating function to positions 52-47 (**Fig. 6b**). In our model, templating and release are flexible at various locations in the template domain, but the main translocation step occurs when the telomeric product is extended to 5’-GGGTT**<u>A</u>**-3’, where A is templated by position 47, rather than 5’-GGTTA**<u>G</u>**-3’, where G is templated by position 46. Re-examining the literature through this lens, we find independent evidence supporting this model. In elegant single-molecule studies, Parks and Stone applied Förster resonance energy transfer (FRET) to investigate template dynamics during the human telomerase catalytic cycle in vitro^41^. RNA-DNA hybrid positioning dynamics were thus observed during elongation of a telomeric primer ending in 5’-…GGG-3’ using specific dNTPs and dideoxy-dNTPs. Interestingly, two distinct positions were observed in equal proportions upon primer extension to 5’-…GGG**TTA**-3’, indicating an equilibrium state of realignment. This result can not be explained under the prevailing model for the telomerase catalytic cycle, but is predicted in our revised model. In the same experiments, upon extension to include the subsequent G residue, yielding 5’-…GGG**TTA<u>G</u>**-3’, there was one major register corresponding that of the re-aligned telomere end. This result was taken as evidence of a shift in equilibrium towards realignment upon synthesis of G templated by position 46C. However, a more likely explanation is that the addition of G templated by position 52 stabilizes the re-aligned state. These orthogonal observations from FRET-based single molecule studies using the wildtype TERC template strongly support our revised model: human telomerase can use a flexible range of templating nucleotides for reverse transcription, but re-alignment readily occurs after incorporation of adenine opposite position 47 of TERC. What underlies this templating base re-positioning relative to *Tetrahymena* and perhaps other telomerase RNAs? Based on phylogenetic and mutagenesis studies in vitro and in human cells, we propose that an alignment domain expansion represented by position 56C during mammalian evolution shifted the dominant templating function of TERC by one nucleotide to positions 52-47. The expanded alignment domain constrains telomerase activity and appears to be dependent on base-pairing or other interactions by cytosine at position 56, in that its deletion or mutation shifts templating function to positions 51-46. Remarkably, by simultaneous mutagenesis of position 56 and position 46C>G, we were able to tag the ends of most chromosomes in human cells with a single C residue. This event was chain terminating and thus double-mutant (56-mutant, 46G) TERC variants could not rescue telomere lengthening or senescence in TERC-null 293T cells. There are thus at least two non-mutually exclusive explanations for the apparent conservation of position 46C, even though it is not a dominant templating residue: 1) it is a vestige of a relatively recent evolutionary event, and 2) its occasional use as a flexible templating base keeps it under evolutionary constraint since introduction of a variant telomeric base at the 3’ end of the telomere impairs further telomere elongation by causing a mismatch with the alignment domain (**Fig. 5g**).

Our study holds several immediate and future implications for the understanding and treatment of human diseases. In families with TBDs, remarkable impacts of TERC template domain mutations have been observed, ranging from complete somatic reversion in hematopoietic stem cells due to their dominant negative effects, to rapid accumulation of variant repeats at telomere ends in successive generations^42,43^. Our results provide a new lens to understand the molecular pathophysiology of TERC variants found in the redefined template region and their consequences on telomere extension in such families. More broadly, the results described here represent the initial insights gained from our ongoing efforts towards an unbiased and comprehensive functional annotation of all TERC variant sequences in a human cellular context, which holds promise to reveal disease mechanisms and inform diagnostics in TBDs. Augmenting TERC is a promising therapeutic strategy for TBDs and other diseases^30,44,45^. In this light, our description here of the first hyperactive TERC variants that can elongate telomeres in TBD patient derived cells is potentially relevant for the development of TERC-based genetic and RNA therapies.

In summary, this study redefines the human telomerase mechanism, based on an unbiased and comprehensive genetic screening of its non-coding RNA template TERC in a cellular context. Effects on native telomere ends were validated through in vitro studies in parallel and iteratively to yield a new functional annotation of the TERC template domain. The model for telomerase activity arising from our findings accounts for previously unexplained observations in independent studies of the telomerase catalytic cycle. Our work offers a robust foundation for re-interpreting existing mechanistic, structural, and genetic data, and guiding future molecular and biophysical investigations of telomerase, which will advance drug development for cancer and aging-associated diseases.

## Methods

### Cell culture

293T cells (ATCC) were grown in DMEM supplemented with 10% fetal bovine serum, MEM Non-Essential Amino Acids, L-glutamate, and penicillin/streptomycin. The TERC-null 293T cell line has been previously described ^29,30^. K562 cells (ATCC) were cultured in RPMI 1640 supplemented with 10% fetal bovine serum, MEM Non-Essential Amino Acids, L-glutamate, and penicillin/streptomycin. Fibroblasts were grown in DMEM supplemented with 15% fetal bovine serum, MEM Non-Essential Amino Acids, L-glutamate, and penicillin/streptomycin. The TBD patient fibroblast line carrying the DKC1 del37L mutation (GM01774^46^) was obtained from the Coriell Cell Repository. The TBD patient fibroblast line carrying the A353V mutation has been previously described^47^. Biological samples were procured under Boston Children’s Hospital Institutional Review Board-approved protocols, after written informed consent in accordance with the Declaration of Helsinki. Both fibroblast lines were stably infected with the pCW57.1 TERT lentivirus. Lentiviral transduction was performed by spinfection at 931G for 2hr in media supplemented with 10µg/ml of protamine sulfate. One day after removal of virus containing media, cells were selected in puromycin at 1µg/ml for 4-7 days. Cell counts were by hemocytometry using trypan blue. Cell images were taken with the Nikon Digital Sight Imager, and white balanced using the ImageJ macro: White balance correction *Version 1.0*.

### Lentivirus Production

Lentiviral production was performed as follows: One ∼70% confluent well of a 6 well plate of 293T cells were transfected with 4.0 µg of transfer plasmid, 1.5 µg of pMD2G, and 2.0 µg PsPax2 using lipofectamine 2000. Two and three days after transfection, the media was harvested, filtered using a 0.45µM filter and stored at -80C. For library scale infections, virus was concentrated by ultracentrifugation at 100,000G for 2 hrs prior to storage at -80C.

### RT-qPCR

Total RNA was isolated using Trizol (Invitrogen) according to the manufacturer’s instructions. RNA was DNAse treated using Turbo DNAse (Invitrogen) according to the manufacturer’s instructions. cDNA synthesis using superscript III reverse transcriptase (Thermo Fisher) with random hexamer priming, and qPCR using ssoAdvanced supermix (Biorad) on a Biorad CFX96 real time PCR machine. Relative gene expression was quantified using ΔΔCt.

### Terminal restriction fragment Southern blotting

DNA was isolated from pelleted cell samples using the PureLink Genomic DNA Mini kit (Invitrogen). 0.5-2.0 µg of gDNA was then digested using RsaI (NEB) and HinfI (NEB) in Cutsmart buffer (NEB) for 2 hours at 37 degrees Celsius. MseI (NEB) was also added to the digestion mix where indicated. Following digestion, samples were re-quantified via nano-drop, and DNA fragments were separated via electrophoresis on a 0.6% agarose gel, followed by Southern blotting onto Hybond-N+ Membrane (Amersham). Membranes were UV crosslinked followed by pre-hybridization for 1 hr using DIG Easy Hyb solution (Roche). Detection of wild type telomere fragments was performed using DIG Easy Hyb supplemented either with the telomeric probe in the TeloTAGGG Telomere Length Assay kit (Roche), or the indicated telomere variant repeat probe, which were synthesized as described ^48^ using the variant probe templates (See oligonucleotides). Telomere variant repeat probes were used at 10 ng / ml. Hybridization was performed at 42C for the wildtype probe and 53C for the variant probes. After hybridization, membranes were washed twice for 5 minutes with 2xSSC 0.1% SDS at room temperature followed by two washes for 15 minutes with 0.2xSSC 0.1% SDS twice at 50C for the WT probe or 53C for the variant probes. Detection was performed as described in the DIG wash and block buffer set (Roche). Membrane stripping was performed by rinsing membranes with MilliQ water followed by two washes for 15 minutes with 0.2M NaOH 0.1% SDS solution at 37C. Quantification of telomere length performed using the WALTER webtool ^49^.

### Saturating variant analysis library design and production

The TERC saturating variant library was generated in the pLKO.1-U3-TERC-500-Puro vector (see below). Five sublibraries were generated by Twist Biosciences with each containing all possible SNVs within a 90 or 91 bp window of the 451bp TERC gene as follows: Library 1 variable region, bp 1-90; Library 2 variable region, bp 91-180; Library 3 variable region, bp 181-270; Library 4 variable region, bp 271-360; Library 5 variable region, bp 361:451. Library representation was confirmed via PCR amplification using Q5 DNA polymerase (NEB) with the TERC gDNA100 F1 and TERC gDNA100 R1 primers followed by next generation sequencing using the Illumina DNA Prep (M) tagmentation kit and the Nova Seq X Plus sequencer with 150 bp paired end reads. Reads were mapped to the pLKO.1 U3-TERC-500 puro insert region using BWA-mem, and variant abundant analysis was performed using ASMV1.0 ^31^. Lentivirus was generated from the library containing plasmids as described above.

### Saturating variant analysis screening

293T-TERC null cells were plated at 1M cells per well of a 12 well dish and transduced via spinfection with one of the 5 sub-library lentiviruses or WT pLKO.1-U3-TERC-500-Puro lentivirus individually. Lentiviral infections performed as described above in two biological replicates. Sufficient cells were infected to maintain >1000X coverage of library variants throughout the infection and culture process. Transduction efficiency was kept below 15%. Uninfected cells were also cultured alongside as a control. After culture for ∼40 days, cells were pelleted and gDNA was isolated using the Purelink genomic DNA isolation kit (Invitrogen). PCR was performed using the TERC gDNA100 F1 and TERC gDNA100 R1 primers and NEB Q5 polymerase in 25 µl reactions with 250 ng DNA per reaction such that each sub-library amplification was performed on at least 4 µg of DNA. Amplification was performed for 28-32 cycles. PCR products were then purified using SPRI beads (Beckman Coulter) followed by library preparation using the Illumina (M) DNA kit and sequencing on a Nova Seq X Plus sequencer with 150 bp paired end reads. Reads were aligned to the pLKO.1 U3-TERC-500 puro sequence using BWA-mem. Variant quantification was performed using ASMV1.0. The Log_2_(fold change) in variant abundance was calculated for experimental samples compared with the input plasmid library for each replicate. The average Log_2_(fold change) across replicates was then calculated for each intended variant. Where indicated, the average Log_2_(fold change) was averaged across variants for a given base. Plots were generated using Matlab R2024b or using GraphPad Prism 10.

### Western Blotting

Cell pellets were lysed with RIPA buffer (Pierce) supplemented with HALT protease inhibitor cocktail (Thermo Fisher), incubated for 30 minutes on ice, then spun at 18,000G for 10 minutes to pellet insoluble material. Lysates were then combined with 2x Laemmli buffer (Bio-Rad), heated to 95C for 5 minutes, and run on 10% SDS-PAGE gels (Bio-Rad) or any KD gels (Bio-Rad), followed by transfer to PVDF membrane (Bio-Rad) using standard procedures. Rabbit anti pCHK2 (Cell Signaling Technologies C13C1) was used at 1:2000 dilution. Primary antibody was detected using HRP conjugated goat anti-Rabbit IgG secondary antibody from Abcam (Ab6721) at 1:5,000 dilution. Blots were then stripped using Restore PLUS Western Blot Stripping Buffer (Thermo Scientific) followed by detection of beta actin using anti-beta actin antibody directly conjugated to HRP (C4; sc-47778 HRP; Santa Cruz Biotechnology) at 1:1000 dilution. Imaging was performed using Bio-Rad ChemiDoc imager.

### Telomere enriched whole genome sequencing

Genomic DNA was purified using the Purelink genomic DNA isolation kit (Invitrogen). 5 µg of gDNA was then digested with RsaI (NEB), HinfI (NEB), MspI (NEB), and HaeIII (NEB) at 37C for 2 hours in Cutsmart buffer (NEB). DNA was then size selected using 0.5x volumes of SPRI beads (Beckman Coulter). On average, 7% of input DNA was recovered. Following size selection, DNA size distribution was evaluated using agarose gel electrophoresis and tape station, with average size of products ∼600-800 bp. DNA libraries were then prepared using the Illumina DNA Prep (M) Tagmentation kit and then subjected to next generation sequencing on a Nova Seq X Plus instrument with 150 bp reads. Telomeric reads were selected and analyzed following the approach described in Hinchie et al ^43^. Briefly, telomeric reads were defined as those containing at least 12 telomeric repeats (5’-GGTTAG-3’, 5’-GGTTAA-3’, 5’-GGTTAT-3’, 5’-GGTTAC-3’, or the corresponding reverse complements). Subsequent analysis was confined to the first 60bp of each read. For the kmer analysis in fig. S4D-F, reads were further filtered by requiring at least 8 telomeric repeats within the first 60 bp of the read, with telomeric repeats including either WT (5’-GGTTAG-3’) or the specific variant of interest as well as the corresponding reverse complements. In fig S4A, counted tracts ranging from 1-8 repeats were flanked by wildtype telomere repeats as in GGTTAG(**GGTTAB)**_n_GGTTAG, where B=T for 46A or 52A, B=C for 46G or 52G, B=A for 46T or 52T. As the read window examined was 60bp, the full length of tracts longer than 8 repeats could not be quantified and were binned as 9+.

### Telomerase purification

293T TERC-null cells were transfected as described above with pBS-U3-hTR-500 vector or the indicated variant pBS-U3-hTR-500 vector and pCDH-3xFLAG-TERT. Two days after transfection, cells were harvested, washed once in PBS, and lysed in 200µl of TRAPeze 1X CHAPS Lysis Buffer (Roche) supplemented with 1:100 of RNasin Plus (Promega) and 1:100 of HALT Protease Inhibitor Cocktail (Thermo Fisher Scientific). Lysates were incubated at 4C for 30 minutes followed by centrifugation to pellet insoluble material. 25µl of Anti-FLAG M2 magnetic beads (Millipore Sigma) that had been washed 3x with CHAPS lysis buffer was then added to lysates followed by incubation for 2 hours at 4C with constant rotation. Beads were then washed four times with 50mM Tris-HCl (pH 8.0), 50mM KCl, 1mM MgCl2, 30% glycerol. Telomerase was then eluted from beads by addition of 30µl of 50mM Tris-HCl (pH 8.0), 50mM KCl, 1mM MgCl2, 30% glycerol, supplemented to 0.5 mg/ml albumin and 0.75 mg/ml 3xFLAG peptide followed by incubation at 4C for 1 hour. After elution, beads were removed by centrifugation at 10,000G with nano-sep 0.45 µM columns (Pall), followed by aliquoting and storage at -80C.

### Direct Telomerase Assays

20µl reactions were prepared using 6 µl of immunopurified telomerase, 5 µM dNTPs (unless otherwise specified), 0.166 μM [α-32P] dCTP (3000 Ci mmol^-1^, Revvity) or 0.166 μM [α-32P] dATP (3000 Ci mmol^-1^, Revvity) as indicated, 1 µM primer (18-GGG unless otherwise indicated) in a final buffer concentration of 50mM Tris-HCl (pH 8.0), 50mM KCl, 1mM MgCl2, 5mM Beta Mercaptoethanol. Reactions were then incubated for 60 minutes at 30C unless otherwise indicated. Reactions were stopped by addition of 100µl stop buffer (3.6 M ammonium acetate and 10mg/ml glycogen) as well as radiolabeled recovery control oligo (see below). 500 µl ethanol was then added to stopped reactions followed by cooling on dry ice for 45 minutes. Reactions were then pelleted, washed with 1ml of 70% ethanol, dried, and resuspended in 10µl TE buffer. Resuspended products were then combined with 10µl of electrophoresis buffer (0.1× TBE, 50 mM EDTA, 0.01% bromophenol blue, 0.01% xylene cyanol and 93% formamide), denatured at 95C for 5 minutes, then centrifuged to precipitate insoluble material. 7µl of sample was then resolved on a 10% polyacrylamide 7M urea gel. After electrophoresis, gels were dried using the Bio-Rad Gel Air drying system and imaged using phosphorimaging with an Amersham Typhoon 5 Biomolecular Imager.

Recovery control oligos were generated by annealing the C template or A template oligo (see below) to the 15-mer oligo by resuspending in 1x buffer 2 (NEB) heating to 95C, then slowly cooling. The annealed oligos were then filled-in by adding 0.166 μM [α-32P] dATP or 0.166 μM [α-32P] dCTP and klenow polymerase (NEB) followed by incubation at room temperature for 20 minutes and purification using the Oligo Clean and Concentrator Kit (Zymo), to generate a radiolabeled 15 mer oligo.

### Telomerase Overexpression Assay

Telomerase overexpression assay was performed as previously described ^50^. Briefly, TERC-null 293T cells were transfected with 2.08 µg TERC and 0.4 µg TERT vector using Lipofectamine 2000 and Opti-mem according to the manufacturer’s instructions. Cells were then cultured for 48 hours, followed by cell harvest, DNA isolation, and terminal restriction fragment Southern blotting as described above. Quantification was performed as previously described ^50^.

### Phylogenetic Tree

Alignment data were used from the Telomerase database ^33^. Phylogenetic tree generated from the taxonomy database using PhyloT v2 ^51^.

### Telomere end sequencing and analysis

gDNA was isolated using the PureLink Genomic DNA Mini kit (Invitrogen). 200-500ng of gDNA was then incubated with TdT (NEB) and 1mM ATP (NEB) in TdT buffer (NEB) at 37C for 2hrs followed by purification using the Oligo Clean and Concentrator Kit (Zymo). Purified ribotailed DNA was then ligated to miRNA cloning linker (NEB) using T4 RNA Ligase 2 Truncated KQ (NEB) in a 10µl volume with 1x T4 RNA ligase buffer and 10% w/v PEG 8000 overnight at room temperature followed by addition of 40 µl of 1x Cutsmart buffer (NEB) supplemented with HinfI (NEB) and incubation at 37C for 1 hr. Reactions were then purified using the Oligo Clean and Concentrator Kit (Zymo). Products were then PCR amplified using the GGTTAG_Junc_F4 or GGTTAG_Junc_F1 primer and the Universal Linker R Primer using Hot Star Taq polymerase (Qiagen) with 35 cycles of amplification. PCR products were purified using 0.9 volumes of SPRI beads (Beckman Coulter) followed by NGS. Reads were filtered for those with positions 20 through 50 of the read with quality scores >= 35. Filtered reads were then searched for the linker sequence, A tails were informatically removed, and the subsequent 6 base pair sequence was quantified for the abundance of permuted telomeric repeats or variant repeats.

### TERC expression vector cloning

The lentiviral TERC expression constructs pLKO.1 U3-TERC-500 puro and pLKO.1 U3-TERC-500 blast were cloned by inserting the U3-TERC-500 insert from the pBS U3-hTR-500 vector into the pLKO.1 puro or blast backbones respectively. TERC variants were cloned into the pBS-U3-hTR-500, pLKO.1 U3-TERC-500 puro, or pLKO.1 U3-TERC-500 blast vectors using the Q5 SDM kit (NEB).

### Plasmids

pLKO.1 puro was a gift from B. Weinberg (plasmid 8453; Addgene)

pLKO.1 blast was a gift from K. Mostov (plasmid 26655; Addgene)

psPAX2 was a gift from D. Trono (plasmid 12260; Addgene)

pMD2.G was a gift from D. Trono (plasmid 12259; Addgene)

pBS U3-hTR-500 was a gift from K. Collins (plasmid 28170; Addgene)

pCDH-3xFLAG-TERT was a gift from S. Artandi (Plasmid 51631; Addgene)

pCW57.1 TERT was a gift from C. Reilly^52^

### Statistical Analysis and Figure Generation

Statistical analysis was performed using GraphPad Prism version 10.6.1. Descriptions of the statistical tests used are present in the figure legends. Figures were generated using GraphPad Prism version 10.6.1 or MATLAB R2024B. Structural images were imaged using generated using Chimera X v1.10.1.

## Data Availability

Unprocessed data and reagents will be available upon request.

## Code Availability

Original code used in this manuscript will be made available upon request.

## Acknowledgements

We thank the patients and families for research participation. We thank Z. Belgacem for technical assistance cloning the lentiviral TERC expression constructs. We thank L. Zon, M. Meyerson, S. Myong, R.C. Lindsley, and C. Reilly for critical input. We acknowledge the next-generation sequencing support provided by S10OD032203 via Tufts University Core Facility Genomics Core. We acknowledge the following funding sources: National Institutes of Health Grant R01DK107716 (SA), F30DK135340 (WM), T32GM007753 (WM), T32GM144273 (WM), Boston Children’s Hospital Translational Research Program (SA), Harvard Stem Cell Institute (WM and SA), Team Telomere (WM and SA), Million Dollar Bike Ride/Penn Medicine Orphan Disease Center (SA), Philanthropic gifts (SA).

## Author Information

## Authors and Affiliations

Boston Children’s Hospital, Dana-Farber Cancer Institute, Harvard Medical School, Boston MA, USA

William Mannherz, Luke Homfeldt, Noah Lampl, Suneet Agarwal

## Contributions

W.M and S.A. conceived the study, designed experiments, acquired funding, supervised the study, and wrote the original manuscript. W.M, L.H, and N.L performed experiments. W.M analyzed the data, performed bioinformatic analyses, and prepared the figures. All authors made edits to the manuscript.

---

**Supplementary Table 1.**
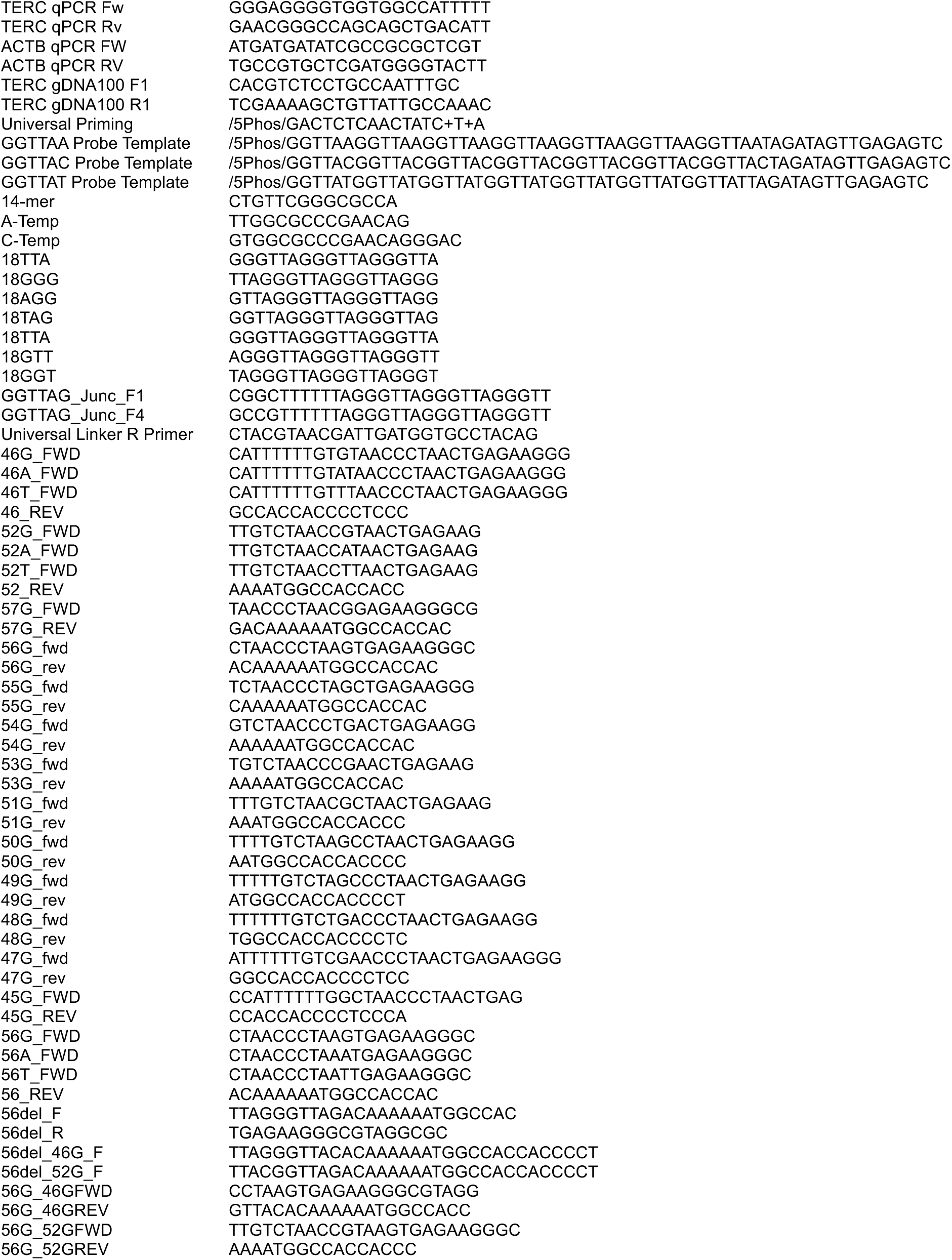
Oligonucleotides. Oligonucleotides synthesized by IDT. Oligonucleotides used in Direct Telomerase Assays were PAGE purified. + indicates locked nucleic acid.

**Extended Data Fig. 1.**
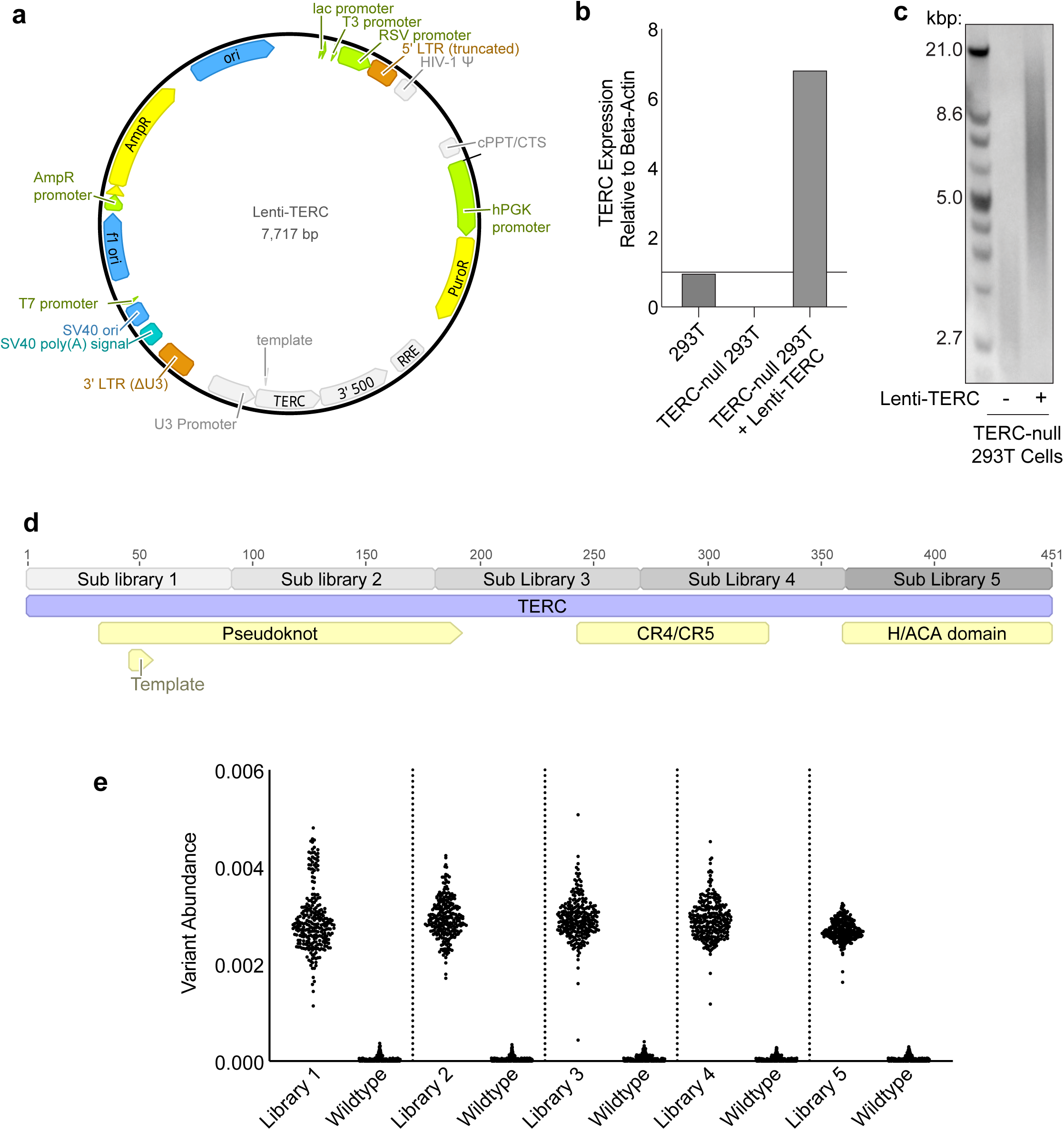
Saturating variant lentiviral libraries of human TERC. **a,** Schematic of lentiviral TERC expression vector. Generated by inserting the U3-TERC-500 insert into the PLKO.1 Puro backbone. **b,** RT-qPCR of TERC expression in the indicated cell lines relative to beta-actin expression. Means of technical triplicates are shown. **c**, TRF Southern blot of TERC-null 293T cells either mock infected or infected with lenti-TERC followed by culture for 17 days. **d,** Schematic of how sub-libraries are divided across the 451 bp TERC gene. The variable regions in libraries 1-4 are 90 bp long. The variable region in library 5 is 91 bp long. **e,** Library inserts from the indicated plasmid DNA was PCR amplified and next generation sequenced followed by quantification of variant abundance (see Methods). Library containing vectors had the intended variants present, while variant abundance in the wildtype vector was minimal. Of note, the variants that were detected in the wildtype vector likely represent PCR or sequencing errors.

**Extended Data Fig. 2.**
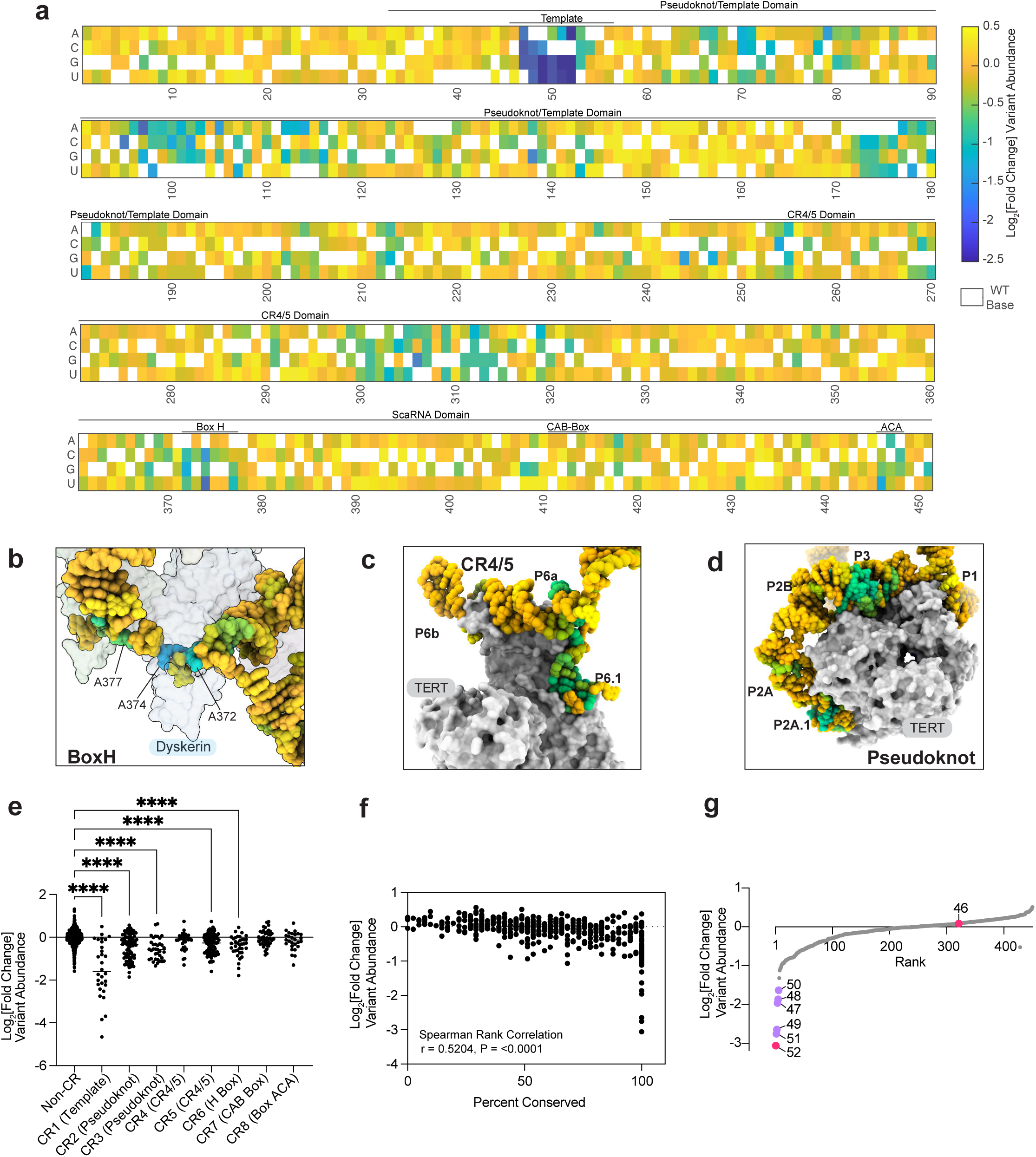
Saturating variant analysis identifies TERC genetic vulnerabilities. **a,** Results from saturating variant analysis of human TERC. Each column represents a single nucleotide. White box indicates the wildtype base. Colored boxes indicate the log_2_[fold change] between a given SNV abundance after culture for 40 days vs plasmid abundance. **b-d**, Base-level results from saturating variant analysis of human TERC. Data represent the average effects of SNVs across a given base pair, overlaid on to the structure of human telomerase. (**b**) PDB: 9QB2^32^; (**c,d**) PDB: 9QAX^32^. Color scale same as in **a**. **e,** Data from saturating variant analysis of TERC broken down by conserved regions (CR). Each dot indicates log_2_[fold change] between a given SNV abundance after culture for 40 days vs plasmid abundance. P values calculated using one way ANOVA with Dunnett’s multiple comparison test. n: non-CR, 963; CR1, 30; CR2, 87; CR3, 36; CR4, 36; CR5, 99; CR6, 36; CR7, 39; CR8, 27. **f,** Data from saturating variant analysis of TERC. Data represent the average effects of SNVs across a given residue plotted against percent that residue is conserved across 42 vertebrates (see methods). P value calculated using Spearman’s Rank Correlation. n=451 **g,** Data from saturating variant analysis of TERC, ranking residues based on their average log_2_[fold change] in variant abundance. All data in this figure represent the average of two biological replicates.

**Extended Data Fig. 3.**
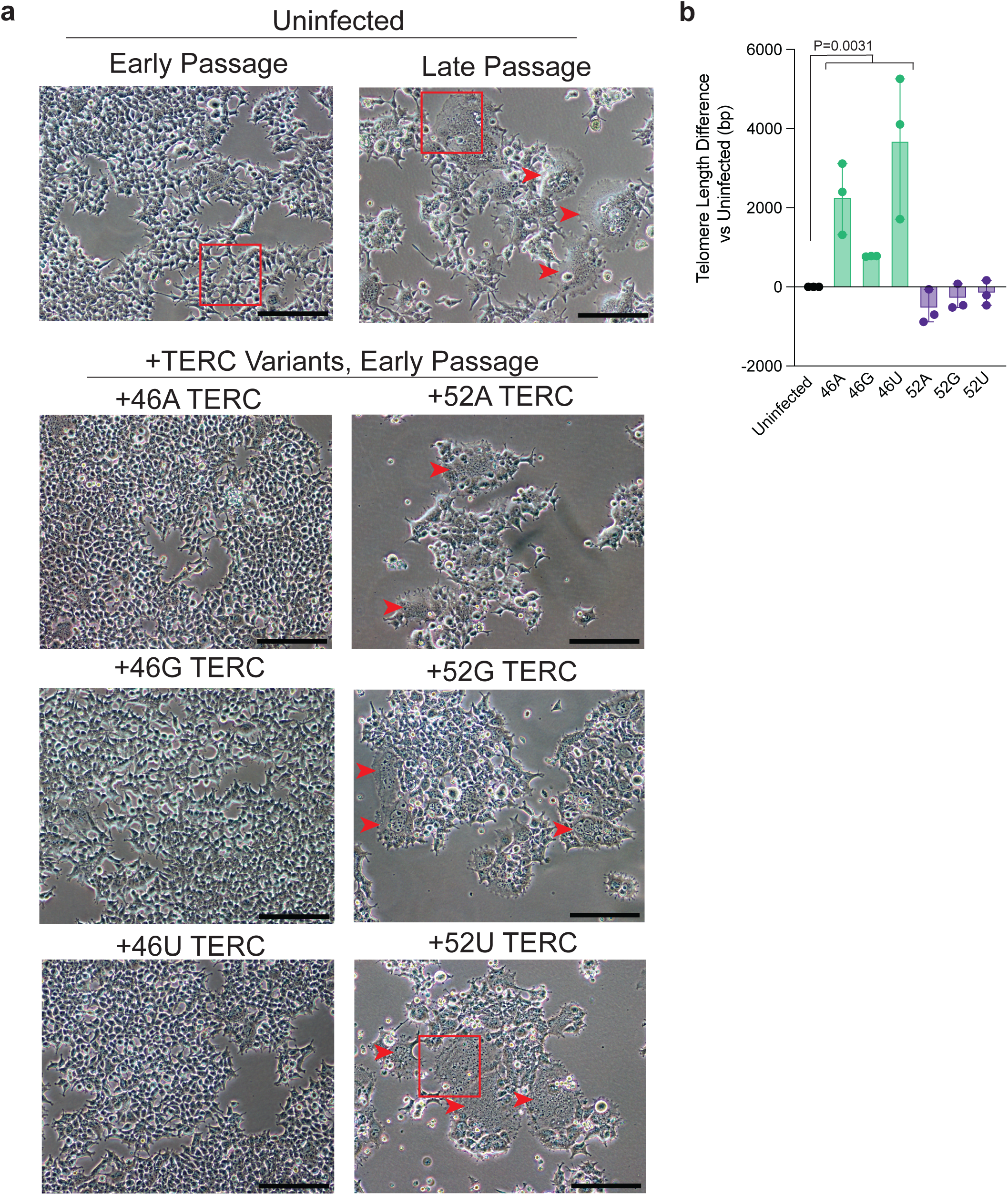
Effects on cellular morphol-ogy and telomere length from position 52 and 46 variant expression support a revised template definition. **a,** Representative images of early passage and late passage TERC-null 293T cells compared with TERC-null 293T cells expressing position 46 or 52 variants. Images of uninfected early passage and TERC expressing cells were obtained 15 days after infection or mock infection. Red arrows highlight areas of syncytia formation. Red squares indicate regions displayed in Fig. 2d. Scale bar is 200µm. **b,** Quantification of mean telomere length WT telomere probing in Fig. 2f. Error bars indicate range. P value calculated for the effect of position 46 variant expression using paired two-sided T test. n=3 biological replicates.

**Extended Data Fig. 4.**
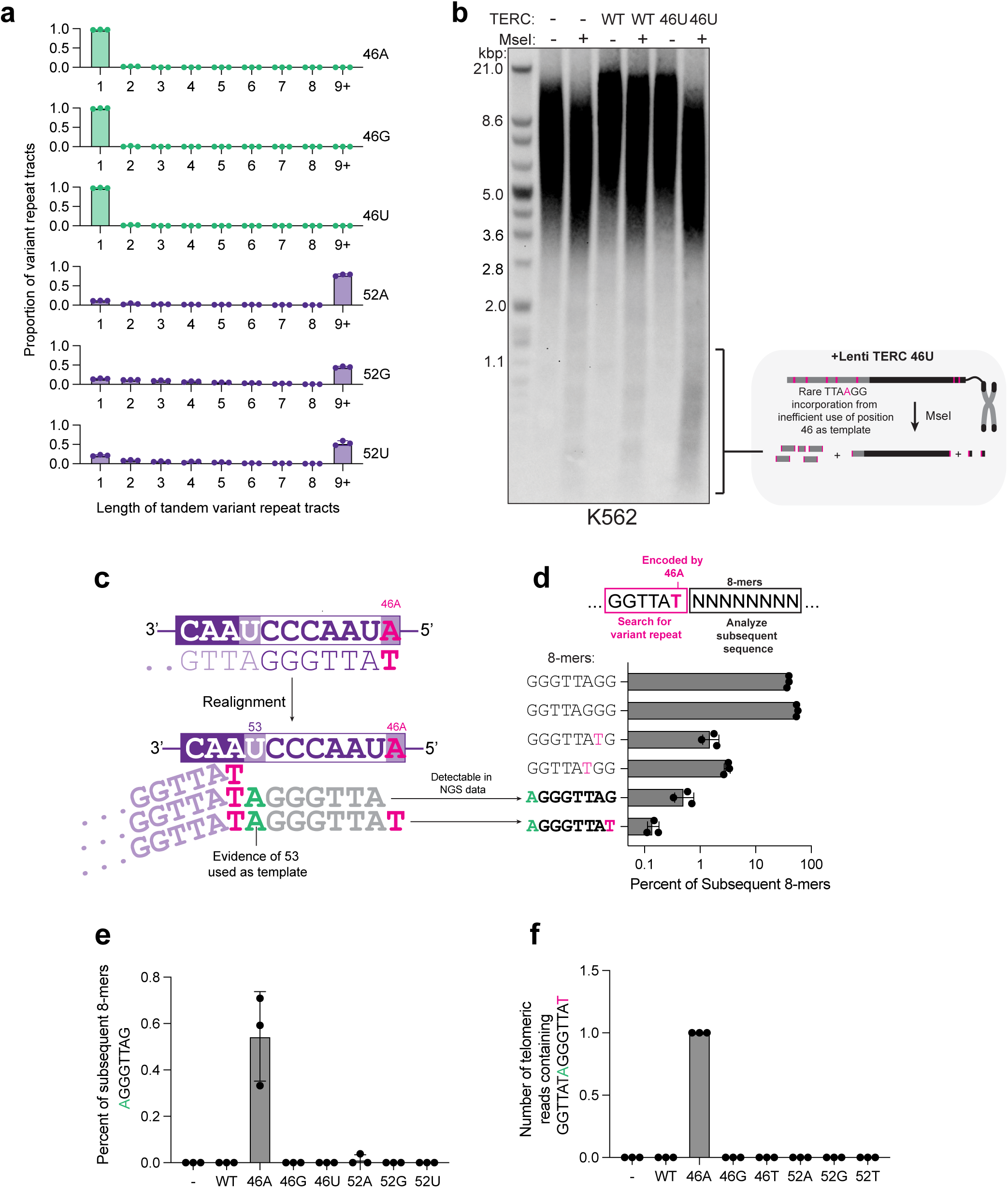
Genomic evidence for flexible use of up to eight TERC templating bases. **a,** Quantification of length of tandem of variant repeat tracts from telomere sequencing dataset (see methods). Results of biological triplicates shown. Error bars indicate SD. **b,** Left, TRF Southern blot of DNA from K562 cells infected with the indicated lentiviral TERC construct and cultured for 19 days performed with or without MseI as indicated. Right, diagram of how MseI digest allows for detection of 46U TERC encoded variants. **c,** Schematic of how 46A usage can be used to mark the end of one telomeric repeat, enabling detection of position 53 templating activity. Synthesis of 5’-**A**GGGTTA**T**-3’ could be consistent with 8nt synthesized in a single extension event. **d,** Quantification of proportion of subsequent 8mers following 5‘-GGTTAT-3’ in 46A TERC expressing cells in telomere sequencing dataset. Error bars indicate SD. Performed in biological triplicate. **e,** Quantification of proportion of 5’-GGTTAT-3’ repeats followed by 5’-**A**GGGTTAG-3’ across samples. Error bars indicate SD. Performed in biological triplicate. **f,** Number of telomeric reads containing the sequence 5’-GGTTAT**A**GGGTTA**T**-3’. Error bars indicate SD. Performed in biological triplicate.

